# miRAW: A deep learning approach to predict miRNA targets by analyzing whole miRNA transcripts

**DOI:** 10.1101/220483

**Authors:** Albert Planas, Xiangfu Zhong, Simon Rayner

## Abstract

**Abstract:** MicroRNAs (miRNAs) are small non-coding RNAs that regulate gene expression by binding to partially complementary regions within the ’UTR of their target genes. Computational methods play an important role in target prediction and assume that the miRNA “seed region” (nt 2 to 8) is required for functional targeting, but typically only identify ∽80% of known bindings. Recent studies have highlighted a role for the entire miRNA, suggesting that a more flexible methodology is needed.

We present a novel approach for miRNA target prediction based on Deep Learning (DL) which, rather than incorporating any knowledge (such as seed regions), investigates the entire miRNA and 3’UTR mRNA nucleotides to learn a uninhibited set of feature descriptors related to the targeting process.

We collected more than 150,000 experimentally validated homo sapiens miRNA:gene targets and cross referenced them with different CLIP-Seq, CLASH and iPAR-CLIP datasets to obtain ∽20,000 validated miRNA:gene exact target sites. Using this data, we implemented and trained a deep neural network - composed of autoencoders and a feed-forward network - able to automatically learn features describing miRNA-mRNA interactions and assess functionality. Predictions were then refined using information such as site location or site accessibility energy.

In a comparison using independent datasets, our DL approach consistently outperformed existing prediction methods, recognizing the seed region as a common feature in the targeting process, but also identifying the role of pairings outside this region. Thermodynamic analysis also suggests that site accessibility plays a role in targeting but that it cannot be used as a sole indicator for functionality. Predictions were then refined using information such as site location or site accessibility energy.

In a comparison using independent datasets, our DL approach consistently outperformed existing prediction methods, recognizing the seed region as a common feature in the targeting process, but also identifying the role of pairings outside this region. Thermodynamic analysis also suggests that site accessibility plays a role in targeting but that it cannot be used as a sole indicator for functionality.

Data and source code available at: https://bitbucket.org/account/user/bipous/projects/MIRAW

**Author summary:** microRNAs are small RNA molecules that regulate biological processes by binding to the 3'UTR of a gene and their dysregulation is associated with several diseases. Computationally predicting these targets remains a challenge as they only partially match their target and so there can be hundreds of targets for a single microRNA. Current tools assume that most of the knowledge defining a microRNA-gene interaction can be captured by analysing the binding produced in the seed region (≈ the first 8nt in the miRNA). However, recent studies show that the whole microRNA can be important and form non-canonical targets. Here, we use a target prediction methodology that relies on deep neural networks to automatically learn the relevant features describing microRNA-gene interactions for predicting microRNA targets. This means we make no assumptions about what is important, leaving the task to the deep neural network. A key part of the work is obtaining a suitable dataset. Thus, we collected and curated more than 150,000 experimentally verified microRNA targets and used them to train the network. Using this approach, we are able to gain a better understanding of non-canonical targets and to improve the accuracy of state-of-the-art prediction tools.

## 1 Introduction

MicroRNAs (miRNAs) are a family of 22-nucleotide (nt) small RNAs that regulate gene expression at the post-transcriptional level. They act by binding to partially complementary sites on target genes to induce cleavage or repression of productive translation, preventing the target gene from producing functional peptides and proteins. Despite advances in understanding miRNA:mRNA interactions, the rules that govern their targeting process are not fully understood [1–4].

While many miRNA targets have been computationally predicted only a limited number have been experimentally validated. Moreover, although a variety of miRNA target prediction algorithms are implemented, results amongst them are generally inconsistent and correctly identifying functional miRNA targets remains a challenging task. The majority of prediction tools are based on the assumption that it is the miRNA seed region – generally defined as a 6 to 8 nucleotide sequence starting at the first or second nucleotide – that contains almost all the important interactions between a miRNA and its target and their focus is on these canonical sites. This seed-centric view has been supported by structural studies [5] and a widely cited report [6] that investigated the importance of other (non-canonical) regions within a miRNA and concluded their contributions had relatively low relevance compared to the (canonical) seed region. However, more recent studies have revealed that many relevant targets are implemented via non-canonical binding and involve nucleotides outside the seed region, indicating that the entire miRNA should be considered in target prediction algorithms [3, 7, 8]. This is also supported by the performance of target prediction tools which typically identify approximately 80% of known miRNA targets, indicating the mechanisms associated with the remaining 20% of non-canonical targets remain poorly understood. Thus, there is an opportunity for novel approaches to improve knowledge of miRNA-regulated processes. In turn, this can lead to better understanding the effects of mutations in the non-coding region of the genome in terms of function and disease. To this end, in this work, we apply deep learning techniques to investigate the role of non-canonical sites and pairing beyond the canonical seed region in microRNA targets.

Almost all target prediction methods are rule-based or adopt machine learning (ML) methodology with varying success. Rule-based systems incorporate various human-crafted descriptors to represent miRNA:gene target binding (e.g. type of pairs in the site, binding stability, or conservation of the target site among species). Machine learning techniques use human crafted descriptors, but as input features to machine learning models. The limitation of both these approaches is the process of feature selection and representation, which is constrained by the use of handcrafted descriptors to model a process that is not fully understood.

Recent increases in computational power have permitted the rise of methods that can directly deal with raw data and autonomously learn and identify patterns to appropriately represent data. In particular, deep learning (DL) [9] has been shown to be an effective method for classification tasks in domains with complex feature representation. Generally, DL methods represent raw data by incorporating multiple hierarchical levels of abstraction. While this approach is typically applied to standard ML problems such as image classification [10], natural language processing [11] or speech recognition [12], it is now finding use in the life sciences for applications such as RNA splicing prediction [13] and gene expression inference [14, 15]. DL has also been applied to the miRNA target prediction problem. Cheng et al. [16] used convolutional neural networks to analyze matrices of miRNA:site features, but the selected features were still human-crafted descriptors and thus the method faces similar problems as rule-based and ML approaches. A more recent work, DeepTarget [17], relied on recurrent neural networks to identify potential binding sites and assess their functionality. However this work is still oriented to the identification of canonical sites and relies on a limited small data set for the training phase.

In this paper we present miRAW, a novel miRNA target prediction tool that works with raw input data and which makes no assumptions about suitable input descriptors. miRAW uses DL to identify relevant descriptors by analyzing the whole mature miRNA transcript, rather than focusing on the seed region, and is trained and tested against experimentally verified positive and negative datasets. When compared to other state-of-the-art miRNA target prediction tools, miRAW demonstrates a significant improvement in performance, highlighting the importance of considering pairing beyond the seed region. In order to gain a deeper understanding of the characteristics of non-canonical targets, we also investigated the prediction results in terms of model design (i.e., how different configurations affect the type of predictions obtained) and from a biological perspective (i.e., how different classes of predicted target sites varied in terms thermodynamic stability and binding structures). In particular, results reveal (i) many potential functional non-canonical binding structures that are supported by experimentally verified miRNA:mRNA target data and (ii) commonly prioritized features such as site accessibility energy and seed region structure are relevant but not sufficient for discerning between functional and non-functional target sites.

## 2 Materials and methods

In our approach, we sought to minimize the introduction of potential biases in the data representation by working directly with the raw data – i.e., the miRNA and mRNA transcripts – rather than incorporating any human selected feature descriptors. To this end we applied deep artificial neural networks (ANN) theory, taking advantage of two of their fundamental properties: (i) with sufficient data-samples and an adequate number of nodes and hidden layers, an ANN can approximate any mathematical function [18]; and (ii) an ANN has the capacity to automatically learn the relevant features of complex data structures by means of its hidden layers [19]. In the following text, we refer to a target site within the 3’UTR of a gene as a miRNA binding site (MBS), comprising the set of nucleotide sites where partially complementary nucleotides individually form bonds between the miRNA and the target mRNA.

The miRAW pipeline (Fig. 1) for investigating the target potential of a miRNA and the 3’UTR of a query gene can be summarized as follows: A 30nt sliding window with a 5nt step is used to scan the 3’UTR of a gene. For each 30nt fragment, miRAW predicts the stability of the binding between the miRNA and the fragment. If the structure is sufficiently stable, miRAW examines the secondary structure to see whether the extended seed region meets the criteria defined in the candidate site selection method (CSSM). If the criteria are met, the sequence of the entire mature miRNA and the 30nt fragment are binarized and fed into miRAW’s neural network to generate a classification. The prediction can be further refined by including one or more filtering steps that apply additional information that is external to the miRNA:site duplex.

**Fig 1.**
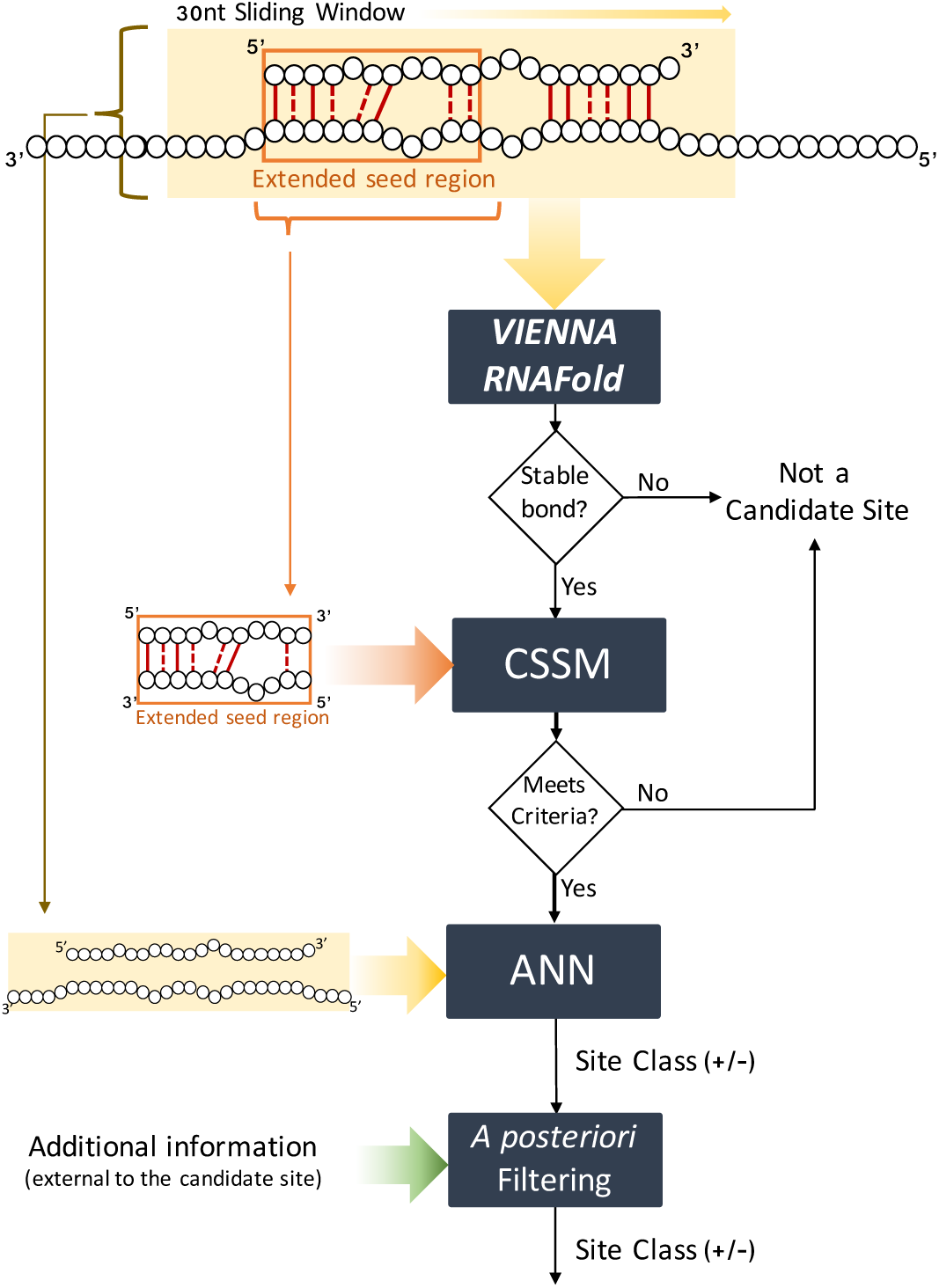
Schematic of the process used by miRAW to evaluate a miRNA Binding Site. (i) A 30nt sliding window is used to scan the 3’UTR of a gene; (ii) The RNAFold software package is used to estimate whether the microRNA and the 30nt transcript can form a stable bond; (iii) If a stable bond is predicted, miRAW checks if the extended seed region meets the criteria defined in the candidate site selection method (CSSM); (iv) If the criteria are met, the full mature microRNA transcript and 30nt corresponding to the candidate site are fed into miRAW’s neural network to generate a classification; (v) The prediction can be refined by a filtering step that applies additional information that is external to the miRNA:site duplex.

### 2.1 Dataset Preparation

A key factor for successful application of any ML classification technique is access to a sufficiently variable and representative dataset that will generalize a trained model to new and unseen data. For the miRNA target prediction problem, this requires a comprehensive dataset of verified positive and negative targets that encompass both canonical and non-canonical examples. While there are multiple data repositories providing information regarding experimentally validated positive miRNA targets [20–22], there are significantly fewer experimentally verified negative targets. This is not an issue for methods that use rule-based approaches to describe positive matches [6], but it represents a major concern for ML-based approaches that require similar numbers of labeled examples for both classes.

Here, we focused on human data and used (i) Diana TarBase [21] – the most comprehensive publicly available dataset, which contains information for both positive (121,090) and negative (2,940) experimentally verified human miRNA:mRNA interactions – and (ii) MirTarBase [20] – containing 410,000 experimentally verified positive targets – as the knowledge core for our study. Annotation related to transcripts and miRNA binding site locations were obtained by cross-referencing Diana TarBase identifiers with miRBase release 21 [23] and Ensembl release 87 [24] entries. As a preliminary step, the Diana and MirTarBase data were parsed to (i) remove inconsistent entries that were marked both as positive and negative targets – due to contradictory results in different experimental validations – and (ii) combine entries that were validated more than once by different verification methods. This produced a final dataset of 33,912 positive (+) and 1,096 negative (-) interactions. The data was then split into two parts (each consisting of 16,496+ and 548- interactions) for the training and testing stages.

#### 2.1.1 Training Dataset

The training dataset serves the purpose of training and validating the ANN responsible for classifying miRNA target sites between functional (positive targets) and non-functional (negative targets). Thus, the training dataset is composed of miRNA:MBS pairs rather than miRNA:mRNA pairs.

- **Positive Training Dataset** To build the positive training dataset we used the reference transcripts of the mature miRNAs and the target mRNAs and, where possible, the binding sites of the experimentally verified targets. However, binding site information is only available and/or parsable for a limited number of Diana Tarbase’s targets. Thus, in order to obtain specific information regarding binding site locations for the remaining target entries, we cross-referenced Diana Tarbase and TarBase with publicly available datasets containing miRNA:MBS locations obtained through PAR-Clip [2] and CLASH [25] experiments. While CLIP and CLASH data provides information regarding experimentally identified miRNA binding site locations, these sites are not necessarily functional. In order to reduce the probability of including non-functional sites in the positive training dataset we considered MBSs that (i) formed stable duplexes -negative free energy in the predicted secondary structure– according to Vienna RNACofold [26] and (ii) corresponded to a miRNA:gene pair marked as functional in mirTarBase or Diana TarBase. Additionally, we complemented our positive training dataset by including the most probable broadly conserved putative sites predicted by TargetScanHuman 7.1 [6] that matched experimentally validated functional data from Diana Tarbase or mirTarBase. The resulting dataset was composed of both canonical and non-canonical MBSs and comprised a total of 22,278 positive target sites for training and validating the miRAW deep learning network.
- **Negative Training Dataset** The smaller number of negative experimentally validated targets poses a challenge when constructing a representative negative dataset. Some ML-based target prediction tools address this problem by using “mock” miRNA targets which are artificially generated miRNA:MBS sequences that resemble true positive targets but which do not appear in positive miRNA target repositories [17, 27]. However, in our case, this type of strategy can lead to the ANN learning the function used to generate the “mock” data and being trained to discriminate between real data and artificial data rather than discriminating functional and non-functional targets. In addition, there is no guarantee that the generated sequences do not belong to miRNA:gene functional pairs yet to be discovered or validated. Thus, we opted for building a negative dataset based upon experimentally verified data. Any sequence of approximately 22 nt within a mRNA of a negatively validated miRNA:mRNA pair represents a possible negative MBS. However, in practice, most of these sequences are irrelevant as they cannot form a stable bond with a miRNA and including them in the training set would merely introduce noise, unnecessarily increasing the complexity of the problem. To obviate this issue, we only considered negative sites within the 3’ UTR of a mRNA that (i) comprise a region with a maximum length of 30 nucleotides and (ii) where a miRNA has the potential to form a stable bond (The choice of a binding site length greater than the average length of an mRNA allows the presence of of bulges within the MBS). For each experimentally verified negative miRNA:mRNA pair, we used a sliding window of 30 nt along the entire 3’UTR region with a 5nt step. The secondary structure of the *miRNA:MBS* duplex was then predicted using the RNACoFold tool from the ViennaRNA package [26] using default settings for all parameters and was considered to be a potential MBS if it had a negative binding energy. This process resulted in a total of 34,918 negatively validated target sites.

For training and validating the neural network, we followed a 10 fold random-subsampling cross-validation approach using the positive and negative training datasets. We stratified the sampling process to ensure the presence of both positive and negative samples for each miRNA family (miRNAs sharing a common ancestor and which have similar similar sequence and structure [23, 28]) present in the training data. 80% percent of data was used for training, 10% reserved for validation and 10% for testing. For each fold we used the same proportion of positive and negative class instances.

#### 2.1.2 Test Dataset

To evaluate the developed method with independent data we generated a dataset using the *∼*17000 experimentally verified miRNA:gene targets excluded from the training data. Note that, in contrast to the training stage, the goal of the test dataset is to evaluate the whole miRAW methodology and, therefore, the testing data consist of pairs containing the miRNA and the whole gene 3’UTR transcripts, rather than the specific MBSs. Again, these 17000 data points were highly biased towards positive entries in a ratio of 97:3 and this imbalance will impede true evaluation of the trained model – a tool that exclusively predicts positive targets against the full test data would achieve an accuracy of 97%. Thus, a testing dataset was generated with equal numbers of positive and negative targets (548+, 548-) where positive entries were randomly selected. To avoid bias attributable to positive target selection, 10 different randomly sampled datasets were generated and compared. Given the strong sequence similarity between miRNAs of the same miRNA family [29, 30], we excluded miRNA:gene pairs that had equivalent pairs (i.e. a pair consisting of a miRNA of the same family that is targeting the same gene) in the training dataset. In this way, we prevent almost identical data pairs from being present both at the training and testing stages.

### 2.2 Candidate Site Selection

Selection of candidate MBSs in a mRNA is another key step for a miRNA target prediction algorithm as it identifies which regions within the mRNA have the potential to be a target binding site. Most target prediction methods follow a similar approach for candidate selection: they scan the 3’UTR of the gene looking for sites that are partially complementary to the miRNA transcript; if a site fulfills certain criteria, it is considered to be a candidate site and is subjected to further analysis. Candidate site selection methods (CSSMs) that focus on the retrieval of canonical targets only consider those sites that have perfect complementary within the miRNA seed region (nucleotides 2 to 8, see Fig. 2a) and will return the smallest number of predicted targets. Methods willing to accept non-canonical sites have looser restrictions: some accept a limited number of bulges, mismatches or wobble pairs in the seed region whilst others accept such mismatches only if there are compensatory nucleotide pairs outside the seed region (Fig. 2b and c).

**Fig 2.**
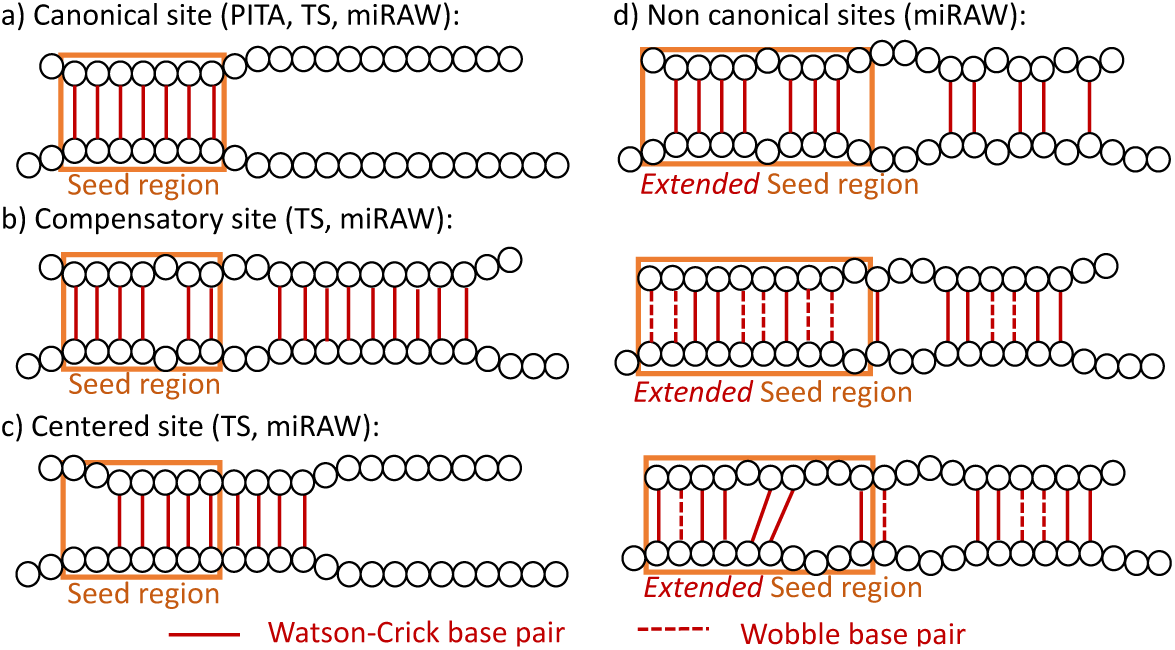
Examples of the types of miRNA binding sites considered by different candidate site selection methods (CSSMs). (a) Potential canonical binding site accepted by the PITA, TargetScan (TS), and miRAW CSSMs. Here, the seed region contains a perfect 7mer. (b) Potential non-canonical compensatory binding site accepted by TS and miRAW CSSMs. The missing nucleotide pair in the seed region is compensated by a 4mer at positions 10 to 14. (c) Potential non-canonical centered target site accepted by TS and miRAW CSSMs. The lack of perfect seed matching is compensated by additional consecutive pairs in nucleotides 9 to 12. (d) Potential non-canonical sites accepted only by the miRAW CSSMs. The extended seed region (10 nucleotides) and the inclusion of wobbles allows these scenarios to be considered as potential target sites.

In an ideal scenario where the training dataset contained sufficient examples of all the possible forms of positive and negative targets, the CSSM would not be required as, theoretically, an ANN would be able to estimate the function acting as CSSM. In reality, there are limited numbers of reliable experimentally verified miRNA:targets (especially for negatively validated sites) and the CSSM step effectively narrows the search space to simplify the ANN classification task.

The CSSM used by miRAW (CSS miRAW) for searching the 3’UTR follows a similar approach to other prediction tools -investigating successive 30-mer segments– but employs a more relaxed set of restrictions that reflect recent experimental studies that relax the requirement of perfect pairing in the seed region and acknowledge a possible role for the other nucleotides. For example, Kim et al [8] report the role of nucleotide 9 in several miRNA binding sites and Grosswendt et al [2] found that a significant number of miRNAs do not require perfect complementarity within the seed region and compensate for this in non-seed nucleotides. Finally, a recent structural study by Klum et al [31] clarify a role for the 3’ end of the miRNA in the targeting process. Based on the findings from these and other related studies, we investigated three different approaches that expand the analysis beyond the typical 7mer seed region and relax the broadly adopted requirement for perfect pairing within the seed region.

In particular, we consider a site to be a candidate MBS if there is a minimum number of base pairs – considering both Watson-Crick (WC) pairing and wobbles – within an extended seed region and investigated three different configurations:

- **CSS miRAW-6-1:10:** a candidate MBS contains at least 6 base pairs between nucleotides 1 and 10.
- **CSS miRAW-7-1:10:** a candidate MBS contains at least 7 base pairs between nucleotides 1 and 10.
- **CSS miRAW-7-2:10:** a candidate MBS contains at least 7 base pairs between nucleotides 2 and 10.

In each case, base pairs do not need to be consecutive in order to accommodate the presence of gaps and bulges.

Thus, these models can accommodate both standard canonical MBSs as well as a broader range of non-canonical target site structures (see Fig. 2), including the vast majority (up to 97.63%) of experimentally validated sites from Diana TarBase and CLIP/CLASH binding site datasets. Moreover, while these relaxed conditions for the seed region generate a much larger number of candidate sites, the decision of whether a site represents a functional target is delegated to the ANN (which considers the entire miRNA & mRNA sequence). In this way, we ensure that minimal assumptions, and hence bias, are incorporated into the analysis.

To further evaluate the impact of choice of CSSM, we also implemented the CSSMs used in two of the most commonly used miRNA target prediction tools:

- **TargetScan (CSS miRAW-TS)** considers three types of sites: (i) perfect canonical matches (perfect complementarity in nt 2 to 8, Fig. 2a), (ii) 3’ compensatory sites (a minimum of 3 consecutive WC pairs between nt 13 and 16 compensates an imperfect seed match -one wobble, bulge or mismatch–, Fig. 2b) and (iii) centered sites (imperfect seed match but 11 contiguous WC pairs between nt 4 and 15, Fig. 2c).
- **PITA (CSS miRAW-Pita)** considers (i) 7mers starting at nt 1 or 2 (Fig. 2a) and (ii) sites containing a gap, wobble or mismatch in the seed region (starting at nt 1) if it contains at least 7 WC pairs.

Both these CSSMs are subsets of CSSM-miRAW-6-1:10 and CSSM-miRAW-7-1:10 (Fig. 2).

Implementation of different CSSMs served a primary purpose of fine-tuning miRAW but also allowed us to investigate the targeting process from a biological perspective. The 5 proposed methods encapsulate different target ranges. At one extreme, CSS-miRAW-TS and CSS-miRAW-P adopt conservative approaches oriented towards canonical sites but they also consider a limited number of non-canonical sites with small irregularities in the seed region; at the other extreme, the other non-canonical CSSMs follow a greedier approach that allows the consideration of several non-canonical sites with broader irregularities in the seed region. These differences produce variations in both the canonical and non-canonical predicted targets.

### 2.3 Transcript Binarization

As an ANN requires numerical data for input, we transformed the miRNA and candidate mRNA site transcripts to binary values using *one hot encoding*. Each of the mRNA and miRNA nucleotides was translated to a binary vector of dimension 4, corresponding to the four possible nucleotide values (see Table 1). Thus, each miRNA target is represented by two concatenated binary vectors: one composed of 120 (4x25nt) dimensions corresponding to the mature miRNA transcript, and a second composed of 160 (4x40nt) dimensions corresponding to the mRNA site (30 nt) and 5 additional upstream and downstream nucleotides.

**Table 1.** Binarized nucleotide encoding

| Nucleotide | Binarization |  |  |  |
| --- | --- | --- | --- | --- |
| A | 0 | 0 | 0 | 1 |
| C | 0 | 0 | 1 | 0 |
| G | 0 | 1 | 0 | 0 |
| U | 1 | 0 | 0 | 0 |
| Empty | 0 | 0 | 0 | 0 |

### 2.4 Neural Network Design

Classification of candidate miRNA:MBSs was performed using a feed forward deep ANN. As we rely on the network to identify the relevant relationships between a sequence and the features that describe the miRNA:mRNA interaction, the input of the network consisted of the binarized transcripts of the miRNA and the MBS. The network was configured so that the number of inputs in the input layer was equal to the dimensionality of the binarized representation of the miRNA:mRNA transcripts, and the output layer consisted of two outputs (positive and negative class classification). In addition, transcripts were aligned so the starting of the seed region corresponded always to the same input node.

The deep ANN was composed of eight dense hidden layers (comprising rectifier activation function -RelU– nodes) whilst the output layer comprised two softmax output nodes. The number of nodes per layer was chosen experimentally using the guidelines in [32] as a starting point and resulted in the structure shown in Supplementary Fig. 1. The first eight hidden layers followed the structure of a stacked autoencoder network and were pre-trained as an autoencoder in order to learn the features that are most representative of miRNA:MBS duplexes (Supplementary Fig. 2). The last three hidden layers and the output layer followed the typical shape of a feedforward classification network. This design was consistent with the functionality of the network: (i) the first hidden layer aims to map the data representation to a higher feature space (ii) the following layers seek relevant features corresponding to the interactions of the inputs (iii) the last two layers are responsible for classifying these features.

To ensure the network’s capacity to deal with newly observed data and to avoid overfitting, training was performed with a dropout rate of 0.2. The maximum number of epochs was set to 500 in order to prevent excessive training time and overfitting. We tested two different loss functions for the network: *negative log likelihood* (NLL) and *cross entropy* (XENT). After performing cross-validation, the trained network that obtained the best performance with its test dataset (XENT Fold 7, see Results) was selected as miRAW’s neural network model.

The two neurons of the output layer correspond to the negative (output 0, *o*_0_) and positive (output 1, *o*_1_) classes. Therefore, the class of the site is determined by the values of the two output neurons:

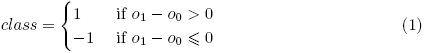

This method will assign a positive or negative classification even if there is only a small difference between the positive and negative output neurons. This scenario corresponds to situations where the network is not confident about the classification of the input data. To deal with such uncertainty a constant parameter *K* was used to define a ‘*grey area*’ in which the network is not able to provide a reliable prediction:

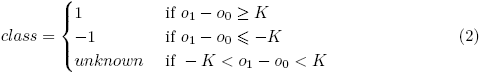

### 2.5 Gene Prediction and Filtering

According to [33], we consider that a miRNA targets an mRNA if any of the potential MBSs of the mRNA are functional. In the representation of the targeting process implemented within miRAW, we require the neural network classify at least one of candidate sites as positive to consider a miRNA:mRNA pair as a positive targeting event.

In our model, given a miRNA *m* and a gene *g*, a candidate site selection method *sm*(*m, g*) determines a set of potential MBSs for that pair, i.e.

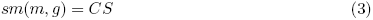

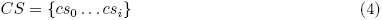

To determine if the miRNA is targeting the gene, each candidate site within the miRNA:mRNA segment is binarized and input to miRAW’s deep ANN. The result of the targeting prediction *T* (*m, g*) corresponds to the disjunction of the neural network outputs (*ann*(*m, cs_i_*)) for all the candidate sites *cs_i_ ∈ CS* in the gene *g*.

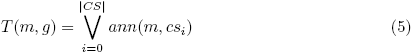

Given that it only requires a single candidate site to be classified as positive for the miRNA:mRNA prediction to be positive means that miRAW is particularly sensitive to false positives. A false negative for a single candidate site can be abrogated by a positive classification for any of the remaining candidate sites but a single false positive cannot be corrected by any number of negative candidate sites. Hence, the more potential sites a CSSM identifies, the higher the probability of obtaining a false-positive prediction.

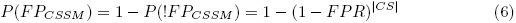

where *F P R* corresponds to the false positive rate of the neural network. This also implies that CSSMs that adopt a greedier approach will end up obtaining more false positives by chance.

The presence of false positives in miRAW’s ANN can be partially attributed to the fact that not all the information concerning miRNA targets can be obtained from the miRNA:MBS duplex and, therefore, cannot be inferred by the neural network. For instance, aspects such as site accessibility [34] require accessing additional external data sources. This external information can be used to refine ANN outcomes by removing sites unlikely to be functional. In an attempt to reduce the likelihood of false positives, we included an *a posteriori* filtering step based on accessibility energy. It is known that miRNA binding sites that are more easily accessible tend to have higher chances of being functional targets [34]; for this reason, several tools usch as PITA, miRMAP [35] or PACMIT [36] combine this information with the binding site minimum free energy (∆*G_duplex_*) to produce a refined target prediction. The site accessibility energy (∆*G_open_*) of a MBS can be defined as the energy required to unfold the secondary structure of the mRNA in order to accommodate the miRNA [34, 36]. As the calculation of ∆*G_open_* requires information that extends beyond the MBS and which involves the whole mRNA sequence, it is particularly well suited for use as *a posteriori* filter in miRAW. Following the site accessibility energy definition of [36], we implemented an ∆*G_open_* filter that removed all predicted sites presenting a ∆*G_open_* higher than a threshold *th_sa_*. Based on results from previous studies [34, 36], we set *th_sa_* = *−*10*kcal/mol*. For accuracy and robustness, we computed local site accessibility following the guidelines defined in [37] and [36]. Specifically, we used the ViennaRNA package [26] and considered the 200nt surrounding the target rather than folding the whole mRNA sequence as this may result in less accurate and more complex secondary structures [36].

### 2.6 Comparison with other miRNA target prediction tools

To assess miRAW’s performance, we compared it against the following commonly used target prediction tools: TargetScan release 7.1 [6], Diana microT-CDS v.4 [38], PITA v.6 [34], miRanda (built upon the mirSVR predictor) [39] and mirDB [40]. These represent the current gold standards (i.e., most commonly referenced) for microRNA target prediction software. These software periodically release datasets containing all predicted miRNA targets using the latest version of the respective tools. In order to evaluate their performance, we cross referenced their latest^1^ available predicted target databases with the unbalanced and balanced datasets defined in Section 2.1. TargetScan offers two different databases in each release, one providing target sites highly conserved among species (TS Conserved) and one providing sites not-necessarily conserved among species (TS NonConserved) and both databases were considered. To assess the significance of the results, we performed a Wilcoxon signed rank test for each of the evaluated metrics; Results were considered significant for *p <* 0.05 unless otherwise stated (see Supplementary Table 4 for specific *p*-values).

### 2.7 Implementation

miRAW was implemented using Java. RNACoFold from the ViennaPackage [26] was used for computing the candidate sites. Implementation of the deep neural network was done using the DeepLearning4Java (DL4J) library [41]. DL4J allows the use of both CPU and GPUs for neural network training and classification. All the analyses presented in this paper were performed using GPUs due to its improved performance; however, a CPU based version of miRAW is also available.

## 3 Results

The two key components of miRAW’s design are (i) the ANN that analyzes candidate target sites and (ii) the CSSM used during the target prediction step. To assess these two aspects of the model we first evaluated the outputs of the ANN training process through cross-validation and then investigated performance using the different candidate site selection methods outlined in the methodology. These comprised the novel (non-canonical) models implemented for miRAW – *miRAW-6-1:10*, *miRAW-7-1:10* and *miRAW-7-2:10* – and the existing (canonical) models already used in TargetScan and PITA – *miRAW-TS* and *miRAW-Pita*. In addition, we explored how *a posteriori* filtering can improve the reliability of the predictions by evaluating miRAW results for the predicted canonical and non-canonical targets in the presence and absence of filtering. Finally, we tested miRAW’s performance by comparing it against TargetScan, Diana microT-CDS, PITA, miRanda and mirDB, which represent the most commonly used target site predictors based on citations.

### 3.1 Neural Network Evaluation

Cross validation of miRAW’s ANN presented good results in terms of predicting both positive and negative sites. This was independent of the loss function used during training, with all evaluated metrics result in scores higher than 0.90 (Fig. 3 and Supplementary Tables 1 and 2). Nonetheless, accuracy and area under the curve (AUC) metrics show that the *XENT* (*accuracy* = 0.92, *AUC* = 0.96) loss function resulted in a statistically significant (Wilcoxon signed sank test) better network compared to the *NLL* function (*accuracy* = 0.91, *AUC* = 0.93). This was reflected in both prediction of positive targets, where the XENT network achieved higher precision, sensitivity and F1-score compared to the NLL network. For negative target prediction, the XENT network returned a larger number of predictions than the NLL but nevertheless achieved a similar negative precision. It is worth noting that, across the different folds, the XENT network was less consistent in terms of negative precision than in positive precision and that for most of the folders it presented more FN than FP. This, combined with the difference in sensitivity and specificity values (0.92 vs 0.94), suggests that the XENT network is slightly biased towards negative predictions as it predicted more negative than positive sites for each fold. Despite this fact, the statistically significant higher accuracy, AUC and F1-scores (both positive and negative) indicate that the XENT network is more appropriate than the NLL network for miRNA target prediction.

**Fig 3.**
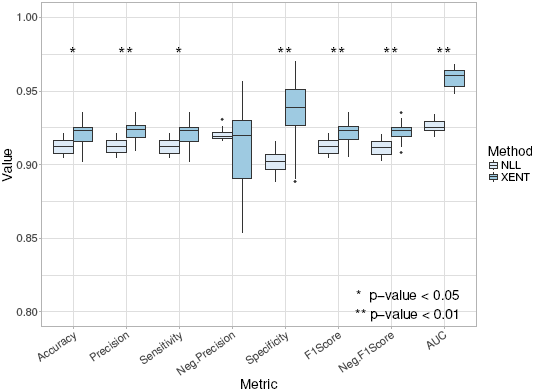
Comparison of miRAW’s neural network performance with the positive and negative training datasets when using a negative log likelihood (NLL) loss function and a cross entropy loss function (XENT) with 10 fold cross validation. XENT provides significantly better accuracy, precision, sensitivity, specificity, F1-scores and area under the curve (AUC) compared to NLL (* p-value*<* 0.05, ** p-value*<* 0.01).

Fig. 4 shows the receiver operating characteristic (ROC) curves for the NLL and XENT networks. The XENT network has a larger AUC, indicating superior performance. Moreover, there is a clear difference in shape of the curves and distribution of data points. The XENT network exhibits a smooth curve with relativity evenly spaced points, the NLL curve is more discontinuous and the data points are concentrated within a smaller region. This indicates a stronger polarization of the NLL network, where all the predictions are strongly classified as a positive or a negative target (i.e. class value is very close to 1 or −1). Conversely, the smoothness of the XENT network represents a more progressive classification, allowing the presence of less polarized predictions, resulting in a more generalized predictive ability. The shape of the NLL curve also suggests that the NLL network might be overfitted and that it might struggle to classify new observations that significantly differ from the training data - this is also supported by the average epoch numbers used by each network to reach its optimal set of weights, 7.32 for the XENT network versus 11.21 for the NLL network.

**Fig 4.**
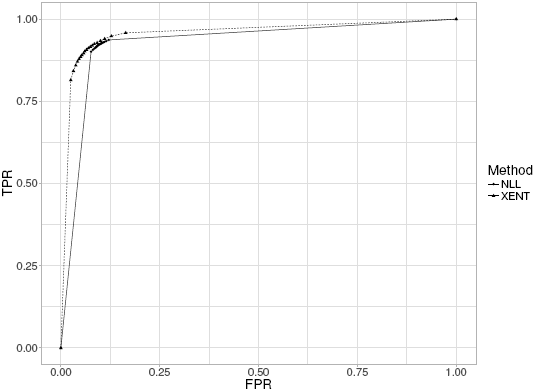
Average ROC Curves for cross validation of miRAW’s neural network using the Positive and Negative training datasets. The dashed line corresponds to the aggregated ROC obtained with the XENT loss function (AUC = 0.96), the solid line corresponds to the NLL loss function (AUC=0.93). The XENT loss function presents a smoother ROC curve with a higher area under the curve, indicating better performance.

The general consistency of calculated parameters and ROC curves across the different folds in the two networks (Supplementary Tables 1 and 2) indicates that the model performance is not dependent on the training and test datasets used. Fold 7 of the XENT network achieved the highest performance in terms of all the considered evaluation metrics and so this ANN model was selected for testing in the gene prediction stage.

### 3.2 miRNA Target Prediction with miRAW: The Role of the Site Selection Method

To investigate the impact of the site selection method, we compared the performance of five different CSSMs (miRAW-6-1:10, miRAW-7-1:10, miRAW-7-2:10, miRAW-TS and miRAW-Pita) in the presence and absence of a site-accessibility energy (AE) filter of −10kcal/mol. The results are summarized in Fig. 5 and Supplementary Table 3. All the methods achieve accuracies between 0.63 and 0.74 with significant differences depending on if the site-accessibility filtering is present (AE) or absent (NF). This effect can be seen when the different CSSMs are ordered by accuracy. miRAW-Pita-NF, miRAW-TS-NF, miRAW-7-2:10-AE and miRAW-7-1:10-AE obtain similar accuracies (*≈* 0.72) with no statistically significant differences in the metric, although miRAW-6-1:10-AE has a slightly poorer performance (0.71). However, miRAW-7-2:10-NF, miRAW-7-1:10-NF, miRAW-Pita-AE and miRAW-TS-NF ranked in the bottom of the table in terms of accuracy. Thus, while the non-canonical CSSMs specifically derived for miRAW obtain better results in the presence of filtering, the canonical derived CSSMs (miRAW-Pita and miRAW-TS) exhibit improved performance in the absence of filtering.

**Fig 5.**
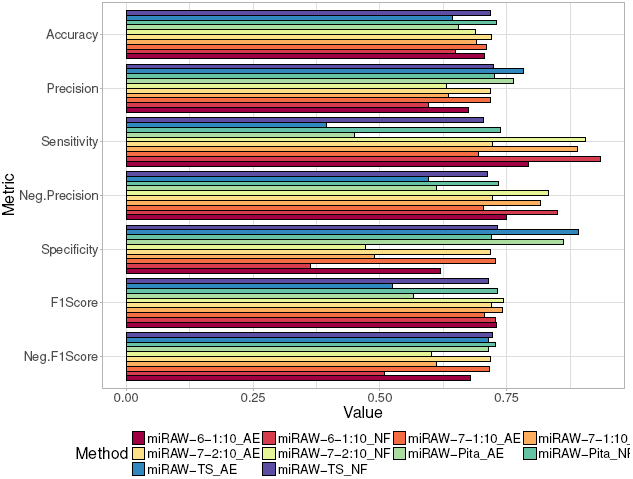
Evaluation of miRAW using different CSSMs and in the presence (AE) and absence (NF) of ∆*G_open_* filtering (threshold = −10Kcal/mol). Results are evaluated in terms of accuracy, precision, sensitivity, negative precision, specificity, positive F1-score and negative F1-score. The best results in terms of accuracy and negative F1-Score were obtained when using Pita’s CSSM and when no filtering was applied. The highest positive F1-Score was obtained by miRAW-7-2:10. Canonical CSSMs (TS and Pita) obtain better results when no filter is applied, the application of ∆*G_open_* filtering introduces false negatives resulting in low sensitivity and negative precision. Conversely, non-canonical CSSMs (miRAW-6-1:10, miRAW-7-1:10 and miRAW-7-2:10) present better results when filtering is applied as this reduces the number of false positives, thereby increasing precision and specificity; when no filtering was applied miRAW was biased towards the prediction of positive sites, which resulted in high sensitivity but low precision.

The F1-scores summarize how well a particular class is classified by a particular CSSM, an optimal CSSM will perform well for both positive and negative targets. Fig. 5 and Supplementary Table 3 show that CSSMs with reported low accuracy underperform in at least one of the F1-scores: miRAW-7-2:10-NF, miRAW-7-1:10-NF and miRAW-6-1:10 have high positive F1-scores but a low negative F1-score caused by an excess of false positives (causing a low specificity) whilst miRAW-Pita-AE and miRAW-TS-AE have a low positive F1-score cause by the excess of false negatives (causing a low sensitivity). However, miRAW-Pita-NF, miRAW-TS-NF, miRAW-7-2:10-AE and miRAW-7-1:10-AE all obtain balanced F1-scores ranging between 0.71 and 0.74, indicating an ability to effectively predict both positive and negative targets.

Fig. 6 summarizes the composition of site types by each CSSM. Fig. 6a shows the average number of canonical (blue) and non-canonical (green) sites identified for each miRNA:gene pair in the test dataset whilst Fig. 6b shows the relative proportions of each identified type. As expected, CSSMs methods following conservative approaches (miRAW-Pita and miRAW-TS) identified more canonical than non-canonical potential sites, whereas the miRAW CSSMs identified larger total numbers of potential sites, of which many more were non-canonical. The figure also shows that the number of predicted canonical sites varies according to the selected CSSM, with the conservative approaches obtaining more canonical sites than the greedy approaches. While this seems to contradict the expectation that all the CSSMs should identify similar canonical sites, the difference can be understood when the higher number of accepted binding structures recognized by the non-canonical oriented CSSMs are taken into account. Several of the sites identified by miRAW-Pita or miRAW-TS overlap with non-canonical binding sites predicted by the miRAW specific CSSMs that present greater stability and are therefore preferentially selected (Fig. 7). Figure 6b also shows that the application of the site accessibility filter does not significantly alter the ratio of canonical and non-canonical sites for any CSSM, suggesting that site accessibility filters do not act as a discriminator between canonical and non-canonical sites.

**Fig 6.**
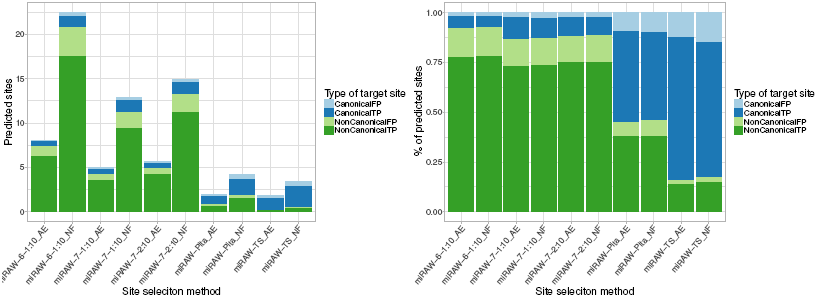
Composition of site types identified by the different CSSMs implemented in miRAW. (a) Average number of miRNA binding sites (MBS) identified by the different CSSMs in a miRNA:mRNA pair. Blue color refer to MBS following a canonical structure, green refer to non-canonical sites; darker colors correspond to positive sites predicted in experimentally verified functional miRNA:mRNA pairs (true positives), lighter colors refer to positive sites identified in non-functional pairs (false positives). (b) Proportion of canonical, non-canonical, true positive and false negative sites identified by each of the candidate site selection methods. Figures illustrate that miRAW-Pita and miRAW-TS CSSMs are strongly biased towards detection of canonical sites whereas miRAW specific CSSMs detect a higher proportion of non-canonical sites.

**Fig 7.**
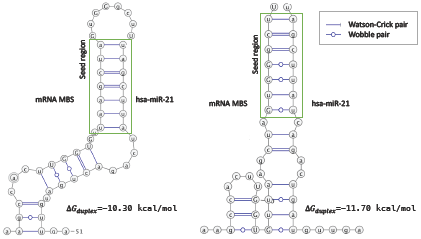
Example of a miRNA binding site that can accommodate a miRNA (hsa-miR-21) with different binding patterns and different site stabilities. The left figure shows a canonical binding (perfect 7mer) with ∆*G_duplex_* = *−*10.30kcal/mol while the right figure shows a non-canonical binding (containing wobbles in the seed region) ∆*G_duplex_* = *−*11.70kcal/mol. While the left structure can be identified by both canonical and non-canonical CSSMs, a non-canonical CSSM will preferentially select the right hand structure as a potential MBS since it reports a more stable predicted binding energy.

### 3.3 Site accessibility filtering

Fig. 5 shows that site accessibility filtering has very different effects in the canonical and non-canonical CSSMs. This difference can be understood by considering the different approaches taken by the canonical (conservative) and non-canonical (greedy) models. The conservative models only consider canonical sites containing close to perfect complementarity (*≥*7mers) in the seed region and a restricted number of non-canonical sites, resulting in a limited amount of candidate sites. Conversely, the greedy models not only recognize canonical sites but also screen a wide range of non-canonical sites that follow unconventional target structures, obtaining a much higher number of potential target sites. This is illustrated in Fig. 8a which shows the average and median number of potential MBSs identified for a miRNA:mRNA pair for each CSSM. The boxplot shows that, in the absence of filtering, more potential MBSs are identified for the more relaxed non-canonical CSSMs. For example, miRAW-Pita NF and miRAW-TS NF identify between 3 and 4 sites respectively per miRNA:mRNA pair, whereas miRAW-7-1:10 NF and miRAW-6-1:10 NF identify 22 and 13 sites respectively. However, as more potential MBSs are identified, the chance of incorrectly obtaining a false positive increases.

**Fig 8.**
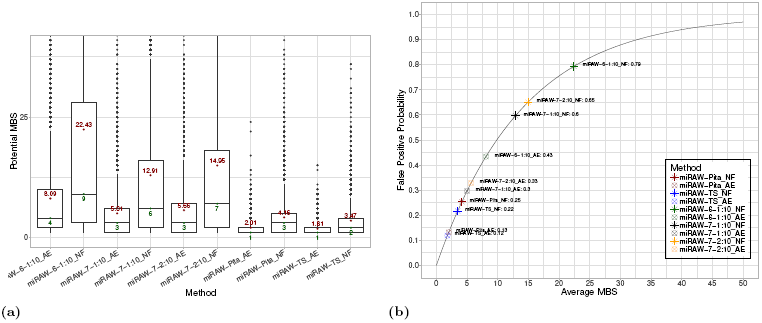
(a)Number of MBSs identified by each CSSM in the presence (AE) and absence (NF) of ∆*G_open_* filtering. Values *>* 40 are excluded from the plot for comparative purposes. Red (upper) numbers and green (lower) numbers show the mean and the median respectively of the number of MBSs identified by each CSSM. miRAW-Pita AE and miRAW-TS AE have the lowest number of MBSs while miRAW 6-1:10 AE has the highest. The number of sites discarded by accessibility energy filtering (AE) is higher in non-canonical oriented CSSMs than in canonical-oriented ones. (b) Relationship between the probability of miRAW obtaining a false positive prediction and the number of sites identified by each CSSM. The fact that miRAW classifies a miRNA:mRNA duplex as positive if a single miRNA:MBSs is predicted as positive by the neural network increases the chances of obtaining a false-positive prediction as the number of potential MBSs increases. As non-canonical oriented CSSMs tend to detect higher numbers of potential MBSs they are more sensitive to a false positive. The application of ∆*G_open_* filtering reduces the number of potential MBSs and therefore reduces the probability of a false positive.

From the training results for the XENT network (Supplementary Table 1), we can estimate the overall probability of the network obtaining a false positive prediction as 0.068 (*P* (*F P*) = 1 *− precision*). However, there is greater variation when we independently consider the various CSSMs. In this case, we can define the probability of obtaining a false positive for a miRNA:mRNA pair using a specific CSSM as:

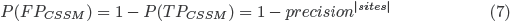

Fig. 8b shows the relationship between the probability of obtaining a false positive and the average number of sites obtained by each CSSM. The non-canonical CSSMs with filtering are at the bottom left of the curve, indicating the efficacy of the filtering step –*a-posteriori* filtering for these CSSMs reduces the number of identified potential MBSs which, in turn, lowers the probability of returning a false-positive error in the network and obtaining a false positive miRNA:mRNA classification–. Conversely, application of *a posteriori* filtering in conservative CSSMs significantly reduces the number of candidate sites, leading to the exclusion of true binding sites and increasing the probability of classifying a miRNA:mRNA pair as a false negative.

We next investigated the relationship between the ∆*G_open_* threshold applied in miRAW and the accuracy and the negative and positive F1-score metrics. The results are summarized in Figs. 9a-c. Again, the canonical and non-canonical CSSMs curves exhibit distinct characteristics. All the metrics for the canonical CSSMs improve with increasing ∆*G_open_*, i.e., these CSSMs are most effective in the absence of any threshold filtering. Conversely, the non-canonical CSSMs improve with low ∆*G_open_* thresholds. While the positive F1-score progressively increases with high ∆*G_open_* thresholds, the negative F1-score achieves a peak value between −8 and −10kcal/mol. This reflects the fact that, as the ∆*G_open_* threshold is increased, more sites are considered by miRAW: increasing sensitivity (more true positives) but in turn reducing specificity (less true negatives and more false positives). The observation that site accessibility filtering increased the performance of non-canonical CSSMs but decreased performance in canonical CSSMs suggests a potential bias towards low site-accessibility energy in non-canonical sites. To explore this possibility, we examined the site accessibility energy distribution of the sites predicted by the different CSSMs and separated them into canonical and non-canonical sites found in true positive (TP) and false positive (FP) miRNA:mRNA pairs. The results are shown in Fig. 10a-d.

**Fig 9.**
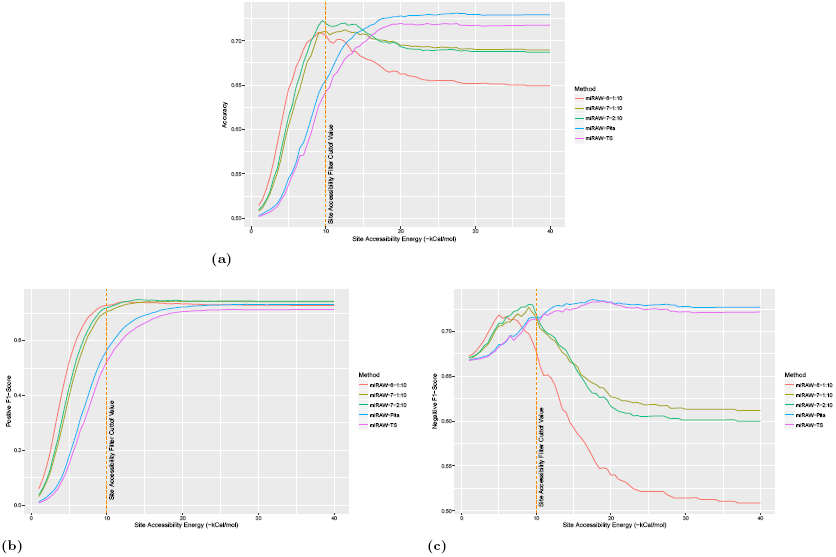
Performance of miRAW in relation to ∆*G_open_* filtering threshold. (a) Variation in accuracy with respect to ∆*G_open_* filtering threshold. (b) Variation in positive F1-score with respect to ∆*G_open_* filtering threshold. (c) Variation in negative F1-score with respect ∆*G_open_* filter threshold. Graphs show that for non-canonical oriented CSSMs, the application of a ∆*G_open_* improves accuracy and negative F1-score values as better scores are obtained when sites with higher ∆*G_open_* values are removed. The peak in the accuracy curve and the fact that the positive F1-score reaches a plateau around ∆*G_open_* = 10, indicates this is an optimal cutoff value. For the canonical-oriented CSSMs, accuracy and positive F1-score metrics reach a plateau around ∆*G_open_ ≥* 23 whereas the negative F1-score curve slightly decreases from ∆*G_open_ ≥* 18. However, the decrease is small compared to the changes in the positive F1-score chart, suggesting that ∆*G_open_* filtering has limited relevance for these models.

**Fig 10.**
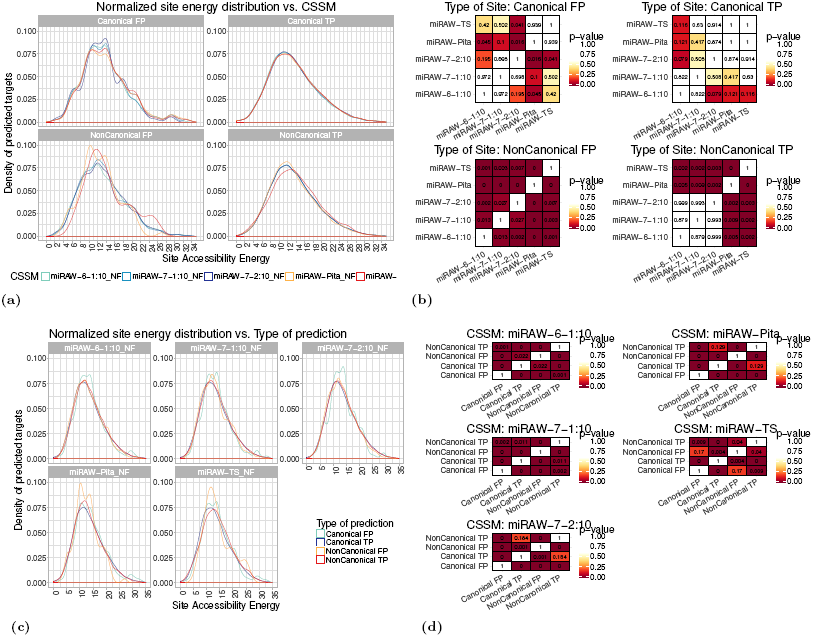
Energy distributions of the site accessibility energy ∆*G_open_* for target sites predicted by miRAW using different CSSMs (a) ∆*G_open_* distributions grouped by the type of site identified by each CSSM (with extreme values removed for comparative purposes). Blue curves correspond to non-canonical CSSMs, red and yellow curves correspond to canonical CSSMs. In general, ∆*G_open_* distributions are smoother for true positive sites (for both canonical and non-canonical CSSMs) than for false positive sites. (b) Pairwise comparison for statistical significance among CSSMs (Kolgomorov-Smirnov test (*p <* 0.05)). The most striking differences are between the ∆*G_open_* distributions of non-canonical false positive sites, with differences identified between all CSSMs. For non-canonical true positive sites, statistical significance is only identified between canonical (miRAW-TS, miRAW-Pita) and the non-canonical (miRAW specific) CSSMs. For canonical sites, there are fewer significant differences; this can be explained by the fact that all the CSSMs identify similar MBSs. (c) Same energy distribution data in (a), but grouped by the CSSMs used for identifying the sites. The smoother distribution of the true positives is also apparent in these plots. (d) Pairwise comparisons of the different site types identified by each CSSM (TP/FP and canonical/non-canonical) - (Kolgomorov-Smirnov test (*p <* 0.05)).

Fig. 10a groups data according to type of site and classification outcome (canonical TP, canonical FP, non-canonical TP and non-canonical FP). For the canonical TP sites, the different CSSM energy distributions do not present statistically significant differences (Kolgomorov-Smirnov test; *p <* 0.05)(Fig. 10b); this is explained by the fact that all the site selection methods identify similar canonical sites. For the FP canonical sites there are no significant differences among the three non-canonical CSSMs, however the miRAW-Pita CSSM does have significant differences with these CSSMs. From the corresponding graph in Fig. 10a (top right) the peak in the miRAW-Pita distribution occurs around a threshold of −10kcal/mol whereas for the non-canonical CSSMs the peaks occur between -14 and -15 kcal/mol. The reason for this difference is unclear as we would anticipate a similar set of false positive canonical sites for all the CSSMs -as all consider sites with perfect seed region complementarity–. However, one possibly for this divergence may be a consequence of the fact that, in contrast to the other CSSMs, miRAW-Pita is the most conservative and does not consider pairing beyond the seed region as a factor for determining the binding site. This is also consistent with the situation shown in Fig. 7 where some canonical sites identified by miRAW-Pita can accommodate more stable non-canonical bindings. This explanation is also consistent with the lower peak in the miRAW-TS CSSM curve, miRAW-TS only considers orthodox non-canonical sites (involving several consecutive WC pairs outside the seed region) compared to miRAW CSSMs. For the non-canonical TPs, there are significant differences between the canonical and non-canonical CSSMs. Although the profiles of the curves for the miRAW-Pita and non-canonical distributions appear similar, there are significantly more sites predicted at the peak energy (*≈*-12kcal/mol) for miRAW-Pita than for the non-canonical CSSMs, although it is unclear whether this is the only feature responsible for the estimated the significant differences between these distributions. Finally, for the non-canonical FP sites, there are significant differences between all the CSSMs, with the non-canonical CSSMs generally presenting distinct distributions compared to the TS and Pita CSSMs. As the exact differences between normalized energy distribution curves were unclear even between statistically significantly different situations, we also performed pairwise comparisons between the mean (Mann-Whitney U test; *p* = 0.05) and median values (Mood’s median test; *p* = 0.05) for each of the distributions. We found that the miRAW-Pita CSSM tended to have higher ∆*G_open_* in non-canonical sites (both TP and FP), which can be attributed to the fact that miRAW-Pita only considers non-canonical sites based on the seed region whereas both TS and CSSM-miRAW accommodate sites beyond this. Therefore MBSs in Pita non-canonical sites have less dependency on accessibility as they only need to accommodate a (smaller) seed region compared to the broader accessibility by the non-canonical sites in the other CSSMs. This is also observed to some degree between miRAW-TS and the miRAW specific CSSMS; miRAW-TS requires several consecutive binding sites outside of the seed region but the miRAW-CSSMs accommodate even more flexible structures.

Fig. 10c shows the same site energy distribution information, but with each plot grouped by CSSM. In all cases, the TP curves have smoother distributions than the FP curves and Kolgomorov-Smirnov tests Figure 10d) report statistically significant differences (*p <* 0.05), between the energy distributions of the predicted canonical and non-canonical sites for almost all the CSSMs. False positives present more irregular distributions in all the CSSMs. Despite these observed differences, there is no clear explanation for either the distinction between canonical or non-canonical sites or true and false positives.

In summary, the results in Fig. 10 indicate that: there are differences in the energy distributions of the sites obtained using different CSSMs; there are differences between canonical and non-canonical sites; and there are differences between the true and false positives energies. Nevertheless, these differences are not sufficient to identify any clear discriminatory features between MBSs, i.e., ∆*G_open_* application of a ∆*G_open_* filter improved performance of CSSMs by reducing the amount of potential MBS (thus reducing the probability of a false positive), rather than by identifying a relationship between ∆*G_open_* and true or false positives.

### 3.4 Comparison with other miRNA Target Predictors

To assess miRAW’s performance, we compared the best configuration of each of the CSSMs – miRAW-Pita-NF, miRAW-TS-NF, miRAW-6-1:10-AE, miRAW-7-1:10-AE and miRAW-7-2:10-AE– against several other target prediction software tools – TargetScan release 7.1 [6], Diana microT-CDS v.4 [38], PITA v.6 [34], miRanda (built upon the mirSVR predictor) [39] and mirDB [40] using the dataset defined in Section 2.1

The results are summarized in Fig. 11 and Supplementary Table 3. All the miRAW implementations generally obtained (statistically significant^2^) better results than each of the other prediction tools for all the evaluated metrics, except for specificity, where most of the methods obtain a similar score, and for precision where mirDB also obtained similar results. Generally, the other methods presented low accuracies: TargetScan, PITA, miRanda and mirDB values were around 0.50; Micro-TDS achieved a value of 0.61, but this was still much less than that reported for miRAW (0.78). For mirDB, miRanda, TS Conserved and TS NonConserved, the low accuracies seem a consequence of their tendency to misclassify true targets as negative; i.e., despite reporting high specificities (*>* 0.70), their negative precision, and hence their F1-score, was low. Conversely, PITA reported better sensitivity than specificity, but obtained similar positive and negative precision. Finally, microT-CDS did not show a particular bias towards any of the classes. It presented balanced specificity (0.63) and sensitivity (0.59) and similar precision (0.62) and negative precision (0.61). Nevertheless, it was still outperformed by all the tested miRAW configurations.

**Fig 11.**
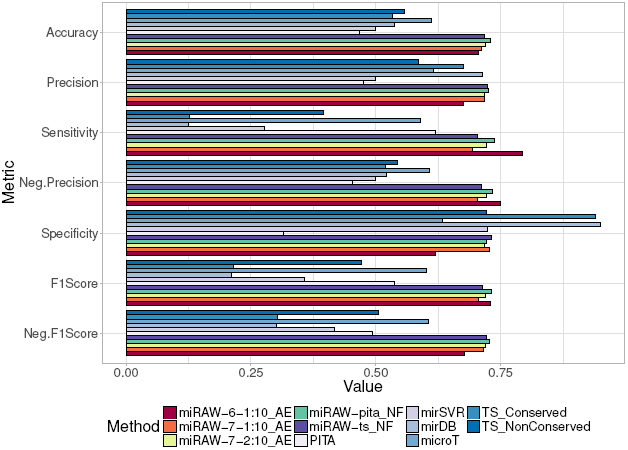
Comparison of miRAW with different CSSMs and six other commonly used target prediction tools (TargetScan C & NC, Diana microT-CDS v4, PITA v6, miRanda and mirDB). Colouring for miRAW results are consistent with the color scheme in Fig. 5; other prediction tools follow a light to dark blue color schema. Evaluation was determined in terms of accuracy, precision, sensitivity, negative precision, specificity and F1-score (an ideal predictor would obtain a score of 1 for each metric). All miRAW configurations outperformed other methods in terms of accuracy and F-scores, which are good representations of general measures of performance. mirDB and Target-Scan (highly conserved targets) obtained high specificity scores but a low negative precision as a consequence of their conservative approach, which classified almost all the miRNA:mRNA pairs as negative. After miRAW, microT was the method which presented better and more balanced results.

These results also highlight how the consideration of interspecies site preservation influences the prediction results. This is particularly apparent in the performance of the two different TargetScan releases. TargetScan achieved low accuracy for both conserved (0.53) and non-conserved datasets (0.56); in both cases the reason for such low accuracy are a consequence of the high number of false negatives. The TS Conserved model presented a high specificity (0.94), meaning that it classified almost all negative targets correctly, but a large number of positive sites were also misclassified as negative, which caused the negative precision to drop to 0.52. Despite filtering positive targets using interspecies conservation information, the TS Conserved precision (0.68) was still lower than the values achieved with any miRAW configuration. For the non-conserved sites dataset (TS NonConserved) the increase in the number of positive predicted sites augmented the number of true positives and sensitivity (0.40) but this, in turn, increased the number of false positives, which reduced precision to 0.59 and specificity to 0.72. Comparison of TargetScan performance with miRAW reveals that both TargetScan releases obtained lower performances than any miRAW configuration in all the evaluated metrics with the exception of specificity, which is a consequence of the conservative approach used by TargetScan regarding positive classification.

Two of the tested methods (PITA an mirSVR) rely on thermodynamic features such as site accessibility ∆*G_open_* or duplex stability ∆*G_duplex_* for target refinement. However, neither method achieved good performance. PITA obtained a relatively high sensitivity compared to other methods (0.62) meaning that it retrieved most of the positive sites, however it has a low precision (0.48) meaning that several negative targets were misclassified. Considering that PITA identifies mostly canonical sites and that it bases its classification on the combination of ∆*G_duplex_* and ∆*G_open_*, this indicates that thermodynamic features are not sufficient for differentiating a positive and a negative target, consistent with our results in (Fig. 8 - Fig. 10). As a consequence several negative targets with low accessibility energy are wrongly classified as positive. Conversely, the miRAW-Pita-NF results, which share the same site selection method, presents better scores in all the evaluated metrics. Considering that miRAW-Pita-NF uses the same canonical target-oriented CSSM as PITA but does not use ∆*G_open_* for determining the functionality of the miRNA:mRNA pair, this also indicates that ∆*G_open_* does not appear to be the most important (i.e. most effective) feature for evaluating canonical sites. This is consistent with the fact that miRAW-Pita-NF and miRAW-TS-NF outperformed miRAW-Pita-AE and miRAW-TS-AE, which only considered sites with low ∆*G_open_*. However, our observation that the methods based on the non-canonical CSSMS had improved performance in the presence of a ∆*G_open_* filter, suggests that this feature does have a role in target functionality. This apparent contradiction can be understood by recognizing that such a role is primarily linked to the broader set of non-canonical sites which correspondingly have a larger range of ∆*G_open_* values, many of which possess higher secondary structure stability, therefore making binding site access difficult.

## 4 Discussion

The imprecise nature of miRNA targeting allows the generation of complex regulatory networks and understanding the mechanisms and functions of these networks requires systematic experimental investigation. In the ideal world it would be possible to experimentally verify the target set of all miRNAs, but both the cost and limited throughput of current methods means that miRNA studies depend on computational predictions to complement experimental data.

The requirement for complimentary base pairing within the seed region for miRNA targeting has been established through numerous experimental studies and forms the basis of all current prediction tools. Initially, it was assumed that seed region binding was a core requirement for all targets but, as more non-canonical targets were experimentally identified, prediction tools evolved to try to accommodate this divergence. The differences in how the various prediction tools recognize the relevance of specific deviations from canonical binding highlights the complexity surrounding the targeting process. The most conservative tools only consider targets that achieve full complementary pairing in the seed regions, whereas other tools allow compensatory binding to accommodate seed mismatches. Moreover, to accommodate non-canonical binding sites, current target prediction tools rely on the use of human crafted descriptors in an attempt to summarize current knowledge regarding miRNA:mRNA interactions, maintaining a bias towards properties associated with the miRNA seed region. Also, as knowledge has increased, so has the complexity of feature descriptors and consequently there is limited consistency amongst the different tools. Thus, researchers tend to adopt a “carpet bombing” approach of target space using multiple prediction tools retaining only those targets that are common among a certain fraction of tools. This further biases predictions back towards the most conservative (i.e. canonical) targets.

In this study, we adopted a neutral approach towards the prediction process, avoiding incorporating any knowledge related to the targeting process. The performance gap between miRAW and the descriptor-based approaches suggests that current knowledge is still not sufficient to accurately capture all aspects of the miRNA targeting process. This is consistent with recent studies, e.g., [3], [7] and [31], which demonstrate that the whole miRNA can play a relevant role in many functional miRNA targets. Based on these findings, we took advantage of deep learning methodology to incorporate the whole miRNA sequence for target prediction. As deep learning has the capacity to automatically extract its own data feature descriptors, miRAW is not limited by the assumptions incorporated into current target prediction tools. Our experiments showed that miRAW consistently outperforms current techniques, suggesting that the descriptors learned by the deep neural network are able to encode current knowledge and include additional yet to be understood information.

Moreover, we attempted to removed any preconceptions from the learning stage by including all miRNA and mRNA nucleotides as input to our model. The only knowledge we apply is during the selection of candidate targets where we implement a selection step to retain miRNA:mRNA pairs that have established binding within a relaxed seed region that spans nucleotides 1 through 10. Despite the application of this selection step, the entire miRNA:mRNA sequence is used as input to the deep learning model. This has the benefit of narrowing the search space while retaining a larger number of candidate targets including includes non-canonical target types. In an ideal scenario, with enough representative positive and negative data samples, the selection step could be skipped as a deep enough neural network should be able to map such information into its weights.

Relaxation of the seed region allows the consideration of both canonical and non-canonical targets, including the ones defined in recent studies that stated the importance of considering nucleotides beyond the 7th position [3, 8]. This also aligns with recent studies which investigated potential binding sites based on microarray expression data that indicate a significant role for miRNA nucleotide 9 [8] and structure studies [31] that demonstrate off-site targeting in the 3’ region of the miRNA is achieved by a pivoting structural element *α* helix-7 within the Ago2 protein that permits rapid making and breaking of miRNA:target base pairs in the 3’ end of the seed region. This allows Ago2 to rapidly screen potential targets to dynamically search for non-canonical sites.

The impact of the candidate site selection model can be seen from the results for different CSSMs within the miRAW model. Conservative approaches (miRAW-Pita and miRAW-TS) presented slightly better accuracies than more relaxed approaches (miRAW-6-1:10,miRAW-7-1:10 and miRAW-7-2:10) but their predictions were heavily biased towards canonical sites. On the other hand, more relaxed models identified a higher number of potential MBSs following both canonical and non-canonical structures. For the latter, the higher number of identified sites generated higher numbers of false positives that decreased precision and specificity. This problem was addressed by post filtering sites with high ∆*G_open_*, increasing accuracy, precision and specificity to levels analogous to the ones obtained by miRAW-Pita and miRAW-TS but with a broader spectrum of binding structures. The contrasting performance between miRAW-Pita, miRAW-TS and the different miRAW specific CSSMs can be understood by the way in which the methods filter the candidate targets. miRAW-Pita has a conservative approach that discards any site containing more than one mismatch within the seed region without considering further the remainder of the mature miRNA transcript. This enhances the reliability of the positive prediction, but at the cost of increasing the number of false negatives as non-canonical sites are discarded. At the other extreme, miRAW adopts a more open strategy to maximize the types of sites (both canonical and non-canonical) that are considered. This more accommodating approach allows the detection of more non-canonical sites, but with the consequence of an increased number of false positives. This argument is also consistent with the results observed for miRAW-TS, which has the most restrictive CSSM - small irregularities are permitted in the seed region but this requires compensatory pairing in the 3’ end of the miRNA. The extended seed region permitted by miRAW leads to selection of positive sites with irregular bindings in the canonical seed region, supporting the argument that pairing beyond the seed region has a more important role. This is observed even with the miRAW-TS and miRAW-Pita CSSMs (which still feed the whole transcript sequences into the machine learning model) which obtain better results than their Pita and TargetScan counterparts.

As deep learning has the capacity to automatically extract its own data feature descriptors, by incorporating the entire miRNA and 3’ UTR target region, miRAW is not limited by the assumptions incorporated into current target prediction tools. Our experiments showed that miRAW consistently outperforms current techniques, suggesting that the descriptors learned by the deep neural network are able to encode current knowledge and include additional yet to be understood information. Furthermore, as the amount of available target data increases, CSSM constraints can be relaxed which, in turn, will facilitate the discovery or disposal of additional non-canonical miRNA binding structures.

Another important task within this work was the processing of different data sources to transform them into suitable training, testing and evaluation datasets. For a ML classifier to learn the patterns needed to distinguish different classes it is necessary not only to have good quality training data but also to have a balanced number of instances for each class. We selected Diana TarBase and mirTarBase as our core data sources as they represented the most comprehensive set of evidence for miRNA:mRNA functional interactions, spanning a range of different experimental methods and providing multiple evidence for many interactions. However, for most of the validated experiments the databases do not provide exact details of the target site for the supported interactions. To obtain reliable binding site information we processed and integrated PAR-CLIP and CLASH datasets -which reveal information regarding binding sites and binding structures but not regarding functionality- and cross-referenced them with TarBase and mirTarBase. Generally, there are many resources for experimentally verified positive data but access to experimentally verified negative data is more scarce. Some approaches solve this problem by generating synthetic negative examples, but these may not accurately represent real negative targets and are particularly inappropriate for the DL approach we implemented here. Thus, we generated our negative data by carefully selecting sites that had both the potential of providing stable miRNA binding and were associated with an experimentally verified negative target.

Despite the enhanced performance demonstrated by miRAW, it is prudent to consider some of the potential limitations of automatic feature learning approaches such as DL. Features learned by a neural network can be difficult to interpret and cannot always be easily mapped into useful knowledge. To address this, studies on neural networks knowledge transferability [42] may aid the interpretation process and is the next logical step in our work. Another issue is incorporating knowledge that is external to the miRNA:MBS duplex transcripts. For example, some of the broadly incorporated features in current prediction tools, such as interspecies conservation or site accessibility energy, cannot be inferred by deep learning as these features are built upon external information not contained in the miRNA:MBS duplex transcript. In miRAW this is addressed by applying *a posteriori* filtering that refines the outcomes of the neural network using external information. As a first test of such *a posteriori* filtering, the ∆*G_open_* filter proved to be effective at narrowing the search space in CSSMs possessing an elevated false positive probability (caused by the high number of identified MBSs). An analysis of the ∆*G_open_* energy distribution showed significant differences between canonical and non-canonical sites, and between true positive and false positive predictions. Nevertheless these differences were not enough to discriminate between these categories, indicating that ∆*G_open_* energy is relevant in the targeting process but is not a sufficient indicator to identify target classes. Regarding interspecies conservation, the fact that miRAW, regardless of the CSSM, outperformed current methods without considering interspecies conservation information suggests that this has limited applicability as a descriptor for miRNA sites. This is also supported by the accuracy and F1-Score results obtained by TargetScan NC, which outperformed the ones obtained by TargetScan C. This is also consistent with research from a recent study which found that interspecies preservation filtering can be disregarded for functionally important non-canonical target sites [2].

Another consideration is the combinatory effect of multiple but weak binding sites which, acting in concert, can have significant functional roles [43]. Consistent with other target prediction tools, miRAW’s binding site centric approach cannot evaluate the joint regulatory effect of multiple potential weak target sites in a mRNA as sites are analyzed independently. Nevertheless, we observe that many miRNA:mRNA duplexes where the CSSM detected a high number of potential MBSs tended to be classified as functional. Although in many cases the classification might simply be a reflection of the increased probability of the ANN producing a false positive, it could also be a consequence of the ANN recognizing features associated with cumulatively strong targeting as proposed by [43]. Similarly, some of the false positive predictions obtained by the non-canonical CSSMs without ∆*_open_* filtering might correspond to weak targets without clear regulatory effects. This hypothesis could be explored in miRAW by updating the ANN in order to add a third class (negative, positive and weak) and by transforming the prediction aggregation equation (5) into a cumulative function. Nevertheless, this approach would require the construction of a training dataset containing enough weak target examples, a challenging task considering that most existing miRNA target resources address the targeting from a binary perspective.

The work presented in this paper focused on the prediction of human miRNA targets, nonetheless the methodology can be readily applied to build target prediction models in other species. Bearing in mind that data availability is crucial for building reliable machine learning classifiers, a logical next step is to implement a target prediction model for mouse. Beyond this, the presented approach will benefit from further experimental studies that will serve to validate new predictions obtained by miRAW but also to generate new experimental data to reliably expand the training of the model. Additionally, considering the *a posteri* filtering step can be applied in retrospective way, it can be used to re-investigate the relevance of some miRNA target descriptors, such as interspecies conservation. Finally, as miRAW considers the whole miRNA:mRNA transcript for its predictions, this also allows the use of miRAW to assess the impact of target site mutations and miRNA isoform variations on the targeting process, which have been shown to have functional roles and characteristic populations that can vary amongst different conditions.

1 January 2017

2 See Supplementary Table 4

## 5 Acknowledgements

The research leading to these results has received funding from the European Union Seventh Framework Program (FP7-PEOPLE-2013-COFUND) under grant agreement n.*^o^* 609020 - Scientia Fellows; and from the Helse Sør Øst project “Integrated Methylation/isomiR/gene Expression (MIG) bio-profiles for prediction of treatment response in rheumatoid arthritis” (Prosjektnummer 2016122). We gratefully acknowledge the support of NVIDIA Corporation with the donation of the GeForce Titan × GPU used for this research.

## References

1. Brennecke J, Stark A, Russell RB, Cohen SM. Principles of microRNA–target recognition. PLoS biol. 2005;3(3):e85.

2. Grosswendt S, Filipchyk A, Manzano M, Klironomos F, Schilling M, Herzog M, et al. Unambiguous identification of miRNA: target site interactions by different types of ligation reactions. Molecular cell. 2014;54(6):1042–1054.

3. Moore MJ, Scheel TK, Luna JM, Park CY, Fak JJ, Nishiuchi E, et al. miRNA-target chimeras reveal miRNA 3 [prime]-end pairing as a major determinant of Argonaute target specificity. Nature communications. 2015;6.

4. Seok H, Ham J, Jang ES, Chi SW. MicroRNA Target Recognition: Insights from Transcriptome-Wide Non-Canonical Interactions. Molecules and cells. 2016;39(5):375.

5. Schirle NT, Sheu-Gruttadauria J, MacRae IJ. Structural basis for microRNA targeting. Science. 2014;346(6209):608–613.

6. Agarwal V, Bell GW, Nam JW, Bartel DP. Predicting effective microRNA target sites in mammalian mRNAs. Elife. 2015;4:e05005.

7. Broughton JP, Lovci MT, Huang JL, Yeo GW, Pasquinelli AE. Pairing beyond the seed supports microRNA targeting specificity. Molecular Cell. 2016;64(2):320–333.

8. Kim D, Sung YM, Park J, Kim S, Kim J, Park J, et al. General rules for functional microRNA targeting. Nature Genetics. 2016;.

9. LeCun Y, Bengio Y, Hinton G. Deep learning. Nature. 2015;521(7553):436–444.

10. Krizhevsky A, Sutskever I, Hinton GE. Imagenet classification with deep convolutional neural networks. In: Advances in neural information processing systems; 2012. p. 10971–1105.

11. Collobert R, Weston J. A unified architecture for natural language processing: Deep neural networks with multitask learning. In: Proceedings of the 25th international conference on Machine learning. ACM; 2008. p. 160–167.

12. Graves A, Mohamed Ar, Hinton G. Speech recognition with deep recurrent neural networks. In: 2013 IEEE international conference on acoustics, speech and signal processing. IEEE; 2013. p. 6645–6649.

13. Alipanahi B, Delong A, Weirauch MT, Frey BJ. Predicting the sequence specificities of DNA-and RNA-binding proteins by deep learning. Nature biotechnology. 2015;.

14. Chen Y, Li Y, Narayan R, Subramanian A, Xie X. Gene expression inference with deep learning. Bioinformatics. 2016; p. btw074.

15. Singh R, Lanchantin J, Robins G, Qi Y. DeepChrome: deep-learning for predicting gene expression from histone modifications. Bioinformatics. 2016;32(17):i639–i648.

16. Cheng S, Guo M, Wang C, Liu X, Liu Y, Wu X. MiRTDL: a deep learning approach for miRNA target prediction. IEEE/ACM Transactions on Computational Biology and Bioinformatics. 2015; p. 1–1. doi: 10.1109/TCBB.2015.2510002.

17. Lee B, Baek J, Park S, Yoon S. deepTarget: end-to-end learning framework for microRNA target prediction using deep recurrent neural networks. arXiv preprint arXiv:160309123. 2016;.

18. Nielsen M. A visual Proof that neural nets can compute any function. In: Artificial Neural Networks and Deep Learning. Determination Press; 2016.

19. Hinton GE, Salakhutdinov RR. Reducing the dimensionality of data with neural networks. Science. 2006;313(5786):504–507.

20. Chou CH, Chang NW, Shrestha S, Hsu SD, Lin YL, Lee WH, et al. miRTarBase 2016: updates to the experimentally validated miRNA-target interactions database. Nucleic acids research. 2016;44(D1):D239–D247.

21. Vlachos IS, Paraskevopoulou MD, Karagkouni D, Georgakilas G, Vergoulis T, Kanellos I, et al. DIANA-TarBase v7. 0: indexing more than half a million experimentally supported miRNA: mRNA interactions. Nucleic acids research. 2015;43(D1):D153–D159.

22. Dweep H, Gretz N. miRWalk2. 0: a comprehensive atlas of microRNA-target interactions. Nature methods. 2015;12(8):697–697.

23. Kozomara A, Griffiths-Jones S. miRBase: annotating high confidence microRNAs using deep sequencing data. Nucleic acids research. 2014;42(D1):D68–D73.

24. Aken BL, Ayling S, Barrell D, Clarke L, Curwen V, Fairley S, et al. The Ensembl gene annotation system. Database. 2016;2016:baw093.

25. Helwak A, Kudla G, Dudnakova T, Tollervey D. Mapping the human miRNA interactome by CLASH reveals frequent noncanonical binding. Cell. 2013;153(3):654–665.

26. Lorenz R, Bernhart SH, Zu Siederdissen CH, Tafer H, Flamm C, Stadler PF, et al. ViennaRNA Package 2.0. Algorithms for Molecular Biology. 2011;6(1):1.

27. Maragkakis M, Alexiou P, Papadopoulos GL, Reczko M, Dalamagas T, Giannopoulos G, et al. Accurate microRNA target prediction correlates with protein repression levels. BMC bioinformatics. 2009;10(1):295.

28. Zou Q, Mao Y, Hu L, Wu Y, Ji Z. miRClassify: an advanced web server for miRNA family classification and annotation. Computers in biology and medicine. 2014;45:157–160.

29. Zou Q, Mao Y, Hu L, Wu Y, Ji Z. miRClassify: an advanced web server for miRNA family classification and annotation. Computers in biology and medicine. 2014;45:157–160.

30. Kamanu TK, Radovanovic A, Archer JA, Bajic VB. Exploration of miRNA families for hypotheses generation. Scientific reports. 2013;3.

31. Klum SM, Chandradoss SD, Schirle NT, Joo C, MacRae IJ. Helix-7 in Argonaute2 shapes the microRNA seed region for rapid target recognition. The EMBO Journal. 2017; p. e201796474.

32. Karsoliya S. Approximating number of hidden layer neurons in multiple hidden layer BPNN architecture. International Journal of Engineering Trends and Technology. 2012;3(6):713–717.

33. Lai EC. Predicting and validating microRNA targets. Genome biology. 2004;5(9):1.

34. Kertesz M, Iovino N, Unnerstall U, Gaul U, Segal E. The role of site accessibility in microRNA target recognition. Nature genetics. 2007;39(10):1278–1284.

35. Vejnar CE, Blum M, Zdobnov EM. miRmap web: comprehensive microRNA target prediction online. Nucleic acids research. 2013;41(W1):W165–W168.

36. Mar´ın RM, Van´ıček J. Efficient use of accessibility in microRNA target prediction. Nucleic acids research. 2010;39(1):19–29.

37. Lange SJ, Maticzka D, Möhl M, Gagnon JN, Brown CM, Backofen R. Global or local? Predicting secondary structure and accessibility in mRNAs. Nucleic acids research. 2012;40(12):5215–5226.

38. Paraskevopoulou MD, Georgakilas G, Kostoulas N, Vlachos IS, Vergoulis T, Reczko M, et al. DIANA-microT web server v5. 0: service integration into miRNA functional analysis workflows. Nucleic acids research. 2013;41(W1):W169–W173.

39. Betel D, Koppal A, Agius P, Sander C, Leslie C. Comprehensive modeling of microRNA targets predicts functional non-conserved and non-canonical sites. Genome biology. 2010;11(8):R90.

40. Wong N, Wang X. miRDB: an online resource for microRNA target prediction and functional annotations. Nucleic acids research. 2014; p. gku1104.

41. Deeplearning4j Development T. Deeplearning4j: Open-source distributed deep learning for the JVM. Apache Software Foundation License. 2016;2.

42. Yosinski J, Clune J, Bengio Y, Lipson H. How transferable are features in deep neural networks? In: Advances in neural information processing systems; 2014. p. 3320–3328.

43. Zhao Y, Shen X, Tang T, Wu CI. Weak Regulation of Many Targets Is Cumulatively Powerful—An Evolutionary Perspective on microRNA Functionality. Molecular Biology and Evolution. 2017; p. msx260. doi:10.1093/molbev/msx260.

